# Transcription-independent TFIIIC-bound sites cluster near heterochromatin boundaries within lamina-associated domains in *C. elegans*

**DOI:** 10.1101/813642

**Authors:** Alexis V. Stutzman, April S. Liang, Vera Beilinson, Kohta Ikegami

## Abstract

**BACKGROUND:** Chromatin organization is central to precise control of gene expression. In various eukaryotic spieces, domains of pervasive *cis*-chromatin interactions demarcate functional domains of the genomes. In nematode *C. elegans*, however, pervasive chromatin contact domains are limited to the dosage-compensated sex chromosome, leaving the principle of *C. elegans* chromatin organization unclear. Transcription Factor III C (TFIIIC) is a basal transcription factor complex for RNA Polymerase III, and is implicated in chromatin organization. TFIIIC binding without RNA Polymerase III co-occupancy, referred to as extra-TFIIIC binding, has been implicated in insulating active and inactive chromatin domains in yeasts, flies, and mammalian cells. Whether extra-TFIIIC sites are present and contribute to chromatin organization in *C. elegans* remains unknown.

**RESULTS:** We identified 504 TFIIIC-bound sites absent of RNA Polymerase III and TATA-binding protein co-occupancy characteristic of extra-TFIIIC sites in *C. elegans* embryos. Extra-TFIIIC sites constituted half of all identified TFIIIC binding sites in the genome. Extra-TFIIIC sites formed dense clusters in *cis*. The clusters of extra-TFIIIC sites were highly over-represented within the distal arm domains of the autosomes that presented a high level of heterochromatin-associated histone H3K9 trimethylation (H3K9me3). Furthermore, extra-TFIIIC clusters were embedded in the lamina-associated domains. Despite the heterochromatin environment of extra-TFIIIC sites, the individual clusters of extra-TFIIIC sites were devoid of and resided near the individual H3K9me3-marked regions.

**CONCLUSION:** Clusters of extra-TFIIIC sites were pervasive in the arm domains of *C. elegans* autosomes, near the outer boundaries of H3K9me3-marked regions. Given the reported activity of extra-TFIIIC sites in heterochromatin insulation in yeasts, our observation raised the possibility that TFIIIC may also demarcate heterochromatin in *C. elegans*.

## BACKGROUND

Eukaryotic genomes are organized into domains of various chromatin features including actively transcribed regions, transcription factor-bound regions, and transcriptionally repressed regions [1–4]. Demarcation of chromatin domains is central to precise control and memory of gene expression patterns. Several proteins have been proposed to have activity in demarcating chromatin domains by acting as a physical boundary [5,6], generating nucleosome depleted regions [7], mediating long-range chromatin interactions [8,9], or tethering chromatin to the nuclear periphery [10]. Despite intense studies [11–15], how chromatin domains are demarcated remains poorly understood.

The genome of nematode *Carnorhabditis elegans* has served as a model to study chromatin organization [3,16–18]. The highest level of chromatin organization in *C. elegans* is the chromatin feature that distinguishes between the X chromosome and the autosomes. The X chromosome in *C. elegans* hermaphrodites is organized into large self-interacting domains that have some features shared with Topologically Associated Domains (TADs) seen in other metazoan genomes. [19–22]. The five autosomes, however, lack robust self-interacting domains [22]. Instead, each autosome can be subdivided into three, megabase-wide domains, the left arm, the right arm, and the center [23]. The center domains display a low recombination rate [24,25], a high density of essential genes [26], and low heterochromatin-associated histone modifications [16,27]. The autosome arms are rich in repetitive elements [23] and heterochromatin-associated histone modifications [16,27], and are associated with the nuclear membrane [18,28,29]. Within these generally euchromatic centers and heterochromatic arms lie kilobase-wide regions of various chromatin states including transcriptionally active and inactive regions [3,4]. While condensins, a highly conserved class of architectural proteins [30], define the boundaries of TAD-like self-interacting domains in the X chromosome [22], the contribution of condensins to autosomal chromatin organization is unclear [14,31]. Furthermore, CTCF, another conserved architectural protein central to defining TAD boundaries in vertebrates, is thought to be lost during the *C. elegans* evolution [32]. How chromatin domains and chromatin states in the *C. elegans* autosomes are demarcated remains an area of active investigation [4,28,33,34].

The transcription factor IIIC complex (TFIIIC) is a general transcription factor required for recruitment of the RNA Polymerase III (Pol III) machinery to diverse classes of small non-coding RNA genes [35]. TFIIIC has also been implicated in chromatin insulation [36,37]. TFIIIC binds DNA sequence elements called the Box-A and Box-B motifs [35]. When participating in Pol III-dependent transcription, TFIIIC binding to Box-A and Box-B motifs results in recruitment of transcription factor IIIB complex (TFIIIB), including TATA-Binding Protein (TBP), which then recruits Pol III [35,38]. By mechanisms that remain unknown, however, TFIIIC is also known to bind DNA without further recruitment of TBP and Pol III [37]. These so-called “extra-TFIIIC sites,” or “ETC,” have been identified in various organisms including yeast [39,40], fly [41], mouse [42], and human [43]. In *Saccharomyces cerevisiae* and *Schizosaccharomyces pombe*, extra-TFIIIC sites exhibit chromatin boundary functions both as heterochromatin barriers and insulators to gene activation [40,44,45]. In addition, extra-TFIIIC sites in these yeast species have been observed at the nuclear periphery, suggesting a contribution to spatial organization of chromosomes [40,46]. In fly, mouse, and human genomes, extra-TFIIIC sites were found in close proximity to architectural proteins including CTCF, condensin, and cohesin [41–43,47]. These studies collectively suggest a conserved role for extra-TFIIIC sites in chromatin insulation and chromosome organization. However, whether extra-TFIIIC sites exist in the *C. elegans* genome is unknown.

In this study, we unveiled extra-TFIIIC sites in the *C. elegans* genome. Extra-TFIIIC sites were highly over-represented within a subset of autosome arms that presented a high level of heterochromatin-associated histone H3K9 trimethylation (H3K9me3). Extra-TFIIIC sites formed dense clusters *in cis* and were embedded in the lamina-associated domains. Despite the heterochromatin environment of extra-TFIIIC sites, the individual clusters of extra-TFIIIC sites were devoid of and resided near the boundaries of H3K9me3-marked regions. Our study thus raised the possibility that, like extra-TFIIIC sites in other organisms, *C. elegans* extra-TFIIIC sites may have a role in demarcating chromatin domains.

## RESULTS

### Half of *C. elegans* TFIIIC binding sites lack Pol III co-occupancy

The TFIIIC complex is a general transcription factor required for the assembly of the RNA Polymerase III (Pol III) machinery at small non-coding RNA genes such as tRNA genes (**Fig. 1A**). Extra-TFIIIC sites are TFIIIC-bound sites lacking Pol III co-occupancy, and are implicated in insulating genomic domains and spatially organizing chromosomes [48]. To determine whether the *C. elegans* genome includes extra-TFIIIC sites, we analyzed the ChIP-seq data published in our previous study [49] for TFIIIC subunits, TFTC-3 (human TFIIIC63/GTF3C3 ortholog) and TFTC-5 (human TFIIIC102/GTF3C5 ortholog) (**Fig. 1B**); the Pol III catalytic subunit RPC-1 (human POLR3A ortholog, “Pol III” hereafter); and the TFIIIB component TBP-1 (human TBP ortholog, “TBP” hereafter) in mixed-stage embryos of the wild-type N2 strain *C. elegans*. We identified 1,029 high-confidence TFIIIC-bound sites exhibiting strong and consistent enrichment for both TFTC-3 and TFTC-5 (**Fig. 1C**). tRNA genes were strongly enriched for TFTC-3, TFTC-5, Pol III, and TBP as expected (**Fig. 1D**). We also observed numerous TFIIIC-bound sites with low or no Pol III and TBP enrichment (**Fig. 1D**). Of the 1,029 TFIIIC-bound sites (**Additional file 1**), we identified 504 sites (49%) with no or very low Pol III and TBP enrichment (**Fig. 1E, F**), which we referred to as extra-TFIIIC sites, following the nomenclature in literature [39–43]. We also identified 525 TFIIIC-bound sites with strong Pol III enrichment (51%), which we referred to as Pol III-bound TFIIIC sites (**Fig. 1E, F**).

**FIGURE 1.**
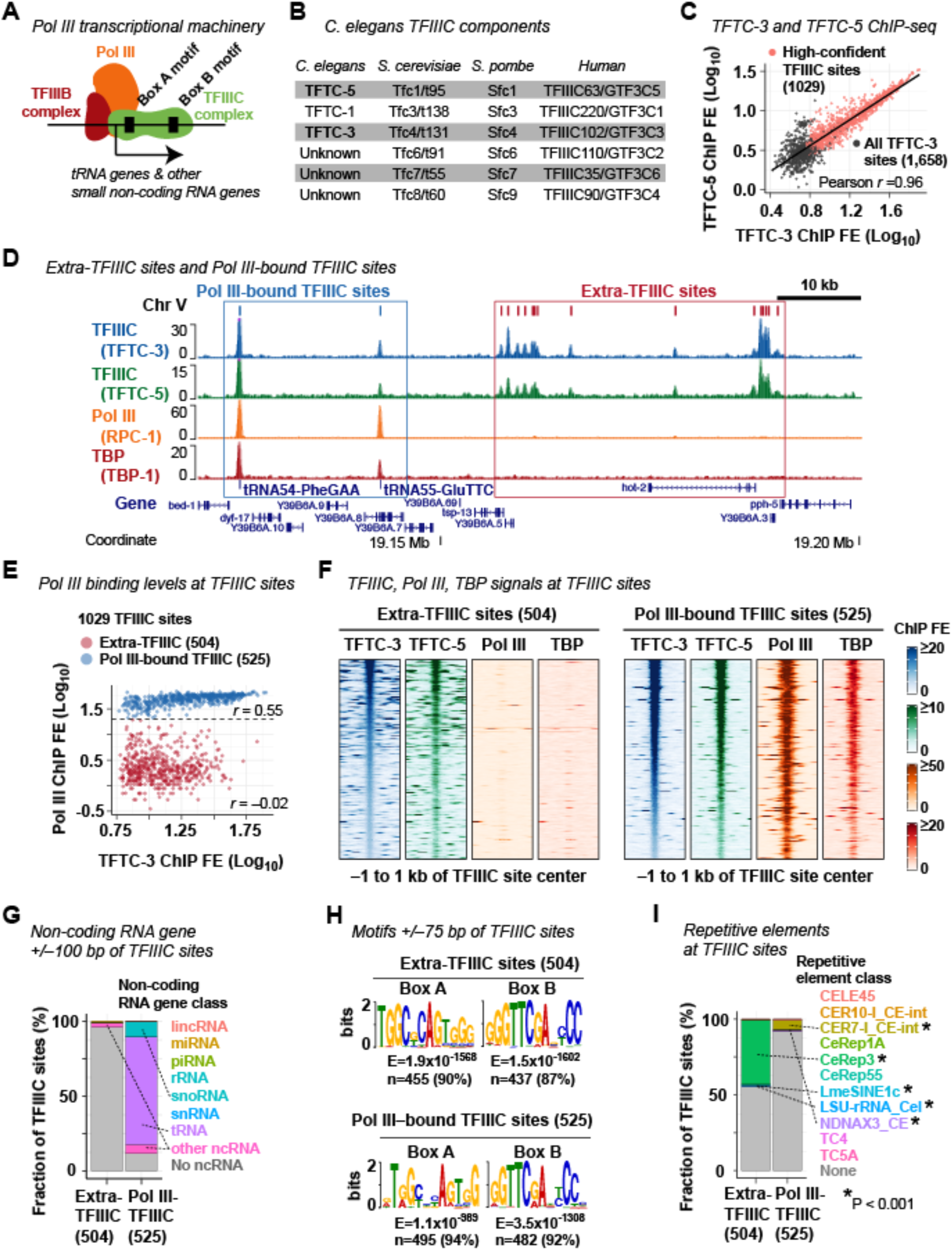
Identification of extra-TFIIIC sites in the *C. elegans* genome. **(A)** A schematic of the RNA polymerase III transcriptional machinery. **(B)** *C. elegans* TFIIIC complex proteins and their yeast and human orthologs for reference. **(C)** Correlation between TFTC-3 and TFTC-5 ChIP-seq fold enrichment (FE) scores at the 1,658 TFTC-3-binding sites. The 1,029 high-confident TFIIIC sites are indicated. **(D)** A representative genomic region showing extra-TFIIIC sites and Pol III-bound TFIIIC sites. **(E)** TFTC-3 and Pol III (RPC-1) FE scores at the 1,029 TFIIIC sites. r, Pearson correlation coefficient within the TFIIIC subclasses. **(F)** TFTC-3, TFTC-5, Pol III (RPC-1), TBP (TBP-1) ChIP-seq FE signals at extra- and Pol III-bound TFIIIC sites. **(G)** Fraction of TFIIIC sites harboring the transcription start site of non-coding RNA genes within ±100 bp of the TFIIIC site center. Dotted lines indicate non-coding RNA gene classes found in ≥10 TFIIIC sites. Pol III-TFIIIC, Pol III-bound TFIIIC sites. **(H)** DNA sequence motifs found in ±75 bp of TFIIIC site centers. **(I)** Fraction of TFIIIC sites overlapping repetitive elements. P, empirical P-value based on 2,000 permutations of the extra- or Pol III-bound TFIIIC sites within chromosomes. Dotted lines indicate repetitive element classes with P < 0.001.

The lack of Pol III and TBP binding in extra-TFIIIC sites may represent a premature Pol III preinitiation complex assembled at Pol III-transcribed non-coding RNA genes [50]. Alternatively, *C. elegans* extra-TFIIIC sites could be unrelated to Pol III transcription and similar to extra-TFIIIC sites reported in other organisms. To distinguish these two possibilities, we examined the presence of transcription start sites (TSSs) of non-coding RNA genes near TFIIIC-bound sites. We observed that only 4% of extra-TFIIIC sites (20/504) harbored TSSs of non-coding RNA genes within 100 bp of the TFIIIC-bound site center (**Fig. 1G**). In contrast, almost all Pol III-bound TFIIIC sites (464 of 525 sites, 88%) harbored TSSs of non-coding RNA genes within 100 bp, and the vast majority of these genes encoded tRNAs (376 sites, 72%) or snoRNAs (52 sites, 10%) (**Fig. 1G**) as expected [35]. Thus, extra-TFIIIC sites in the *C. elegans* genome are unlikely to participate in local Pol III-dependent transcription, a characteristic behavior of extra-TFIIIC sites reported in other organisms [37].

### *C. elegans* extra-TFIIIC sites possess strong Box-A and Box-B motifs

The TFIIIC complex binds the Box-A and Box-B DNA motifs [35] (**Fig. 1A**). However, the majority of extra-TFIIIC sites in yeast and human possess only the Box-B motif [39,40,43]. To determine whether *C. elegans* extra-TFIIIC sites contain Box-A and Box-B motifs, we performed *de novo* DNA motif analyses at extra-TFIIIC sites. Almost all of the 504 extra-TFIIIC sites in *C. elegans* harbored both the Box-A and Box-B motifs (90% with Box-A, E=1.9 x 10^−1568^; 87% with Box-B, E=1.5×10^−1602^; **Fig. 1H**). The pervasiveness of these motifs in extra-TFIIIC sites was comparable to that in Pol III-bound TFIIIC binding sites (94% with Box-A, E=1.1×10^−589^; and 92% with Box-B, E=3.5×10^−1308^) (**Fig. 1H**).

Because the Box-A and Box-B motifs constitute the gene-internal promoter for tRNA genes in eukaryotic genomes [35] (**Fig. 1A**), we hypothesized that extra-TFIIIC sites correspond to genetic elements similar to tRNA genes. In *C. elegans*, a class of interspersed repetitive elements called CeRep3 has been suspected as tRNA pseudogenes [51]. To determine whether extra-TFIIIC sites coincide with repetitive elements, we surveyed the overlap between extra-TFIIIC sites and all annotated repetitive elements (**Fig. 1I**). Strikingly, 44.6% (225 sites) of extra-TFIIIC sites overlapped repetitive elements (permutation-based empirical P<0.001), and almost all of the overlapped elements (95.1%; 214 sites) were the CeRep3 class of repetitive elements (**Fig. 1I**). In contrast, although significant, only 8.2% (43 sites) of Pol III-bound TFIIIC sites overlapped repetitive elements of any class (permutation-based empirical P<0.001). Therefore, unlike extra-TFIIIC sites in yeast and humans, *C. elegans* extra-TFIIIC sites harbored both the Box-A and Box-B motifs; furthermore, a large fraction of these sites corresponded to a class of putative tRNA pseudogenes CeRep3.

### *C. elegans* extra-TFIIIC sites are not associated with regulatory elements for protein-coding genes

Previous studies in human and *S. cerevisiae* reported that extra-TFIIIC sites are overrepresented near protein-coding gene promoters, proposing a role in regulating RNA Polymerase II (Pol II)-dependent transcription [39,43]. To determine whether *C. elegans* extra-TFIIIC sites were located near protein-coding genes, we measured the distance from extra-TFIIIC sites to TSSs of nearest protein-coding genes. *C. elegans* extra-TFIIIC sites were not located near protein-coding gene TSSs compared with Pol III-bound TFIIIC sites (Mann Whitney *U* test, P=0.01) or with randomly permutated extra-TFIIIC sites (Mann-Whitney *U* test, P=2×10^−5^; **Fig. 2A**). Extra-TFIIIC sites were overrepresented in introns and underrepresented in exons and 3′-UTRs, so were Pol III-bound TFIIIC sites (permutation-based empirical P<0.001; **Fig. 2B**). Thus, extra-TFIIIC sites were not differentially overrepresented near protein-coding gene TSSs or in exons, introns, and 3′-UTRs compared with Pol III-bound TFIIIC sites.

**FIGURE 2.**
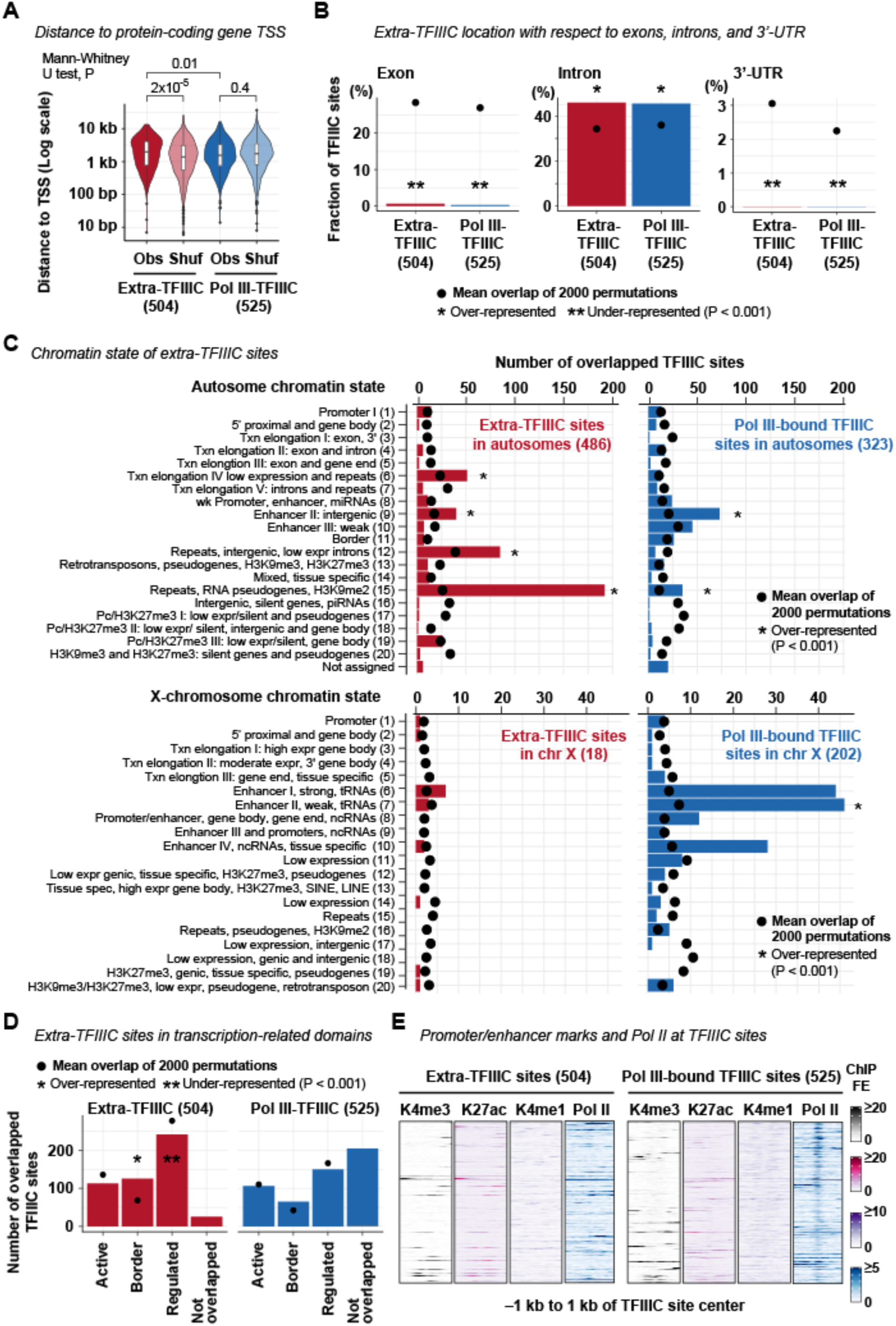
Extra-TFIIIC sites do not reside in protein-coding gene promoters or enhancers. **(A)** Distance between TFIIIC site center and the transcription start site (TSS) of protein-coding genes. Obs, the observed distance. Shuf, the distance of a permutated set of TFIIIC sites (one permutation within chromosomes) to TSS. **(B)** Fraction of TFIIIC sites located in exons, introns, and 3’-UTRs of protein-coding genes. Permutations of TFIIIC sites were performed within chromosomal domains (i.e. center and left/right arms). P, empirical P-value based on the 2000 permutations. **(C)** Chromatin state annotation of TFIIIC sites. The chromatin state annotation reported by Evans et al. (2016) is used. Permutations and P-value computation are performed as in **(B). (D)** Number of TFIIIC sites resided in the three transcription-related domains reported by Evans et al. (2016). Permutations and P-value computation are performed as in **(B). (D)** H3K4me3, H3K27ac, H3K4me1, and RNA Polymerase II (Pol II) fold-enrichment scores (FE) at TFIIIC sites.

To further investigate the relationship between extra-TFIIIC sites and *cis-*regulatory elements for protein-coding genes, we examined the chromatin states defined by a combination of histone modifications in early embryos [4]. Extra-TFIIIC sites were not overrepresented within “promoter” regions (14 sites, 2.8%, permutation-based empirical P=0.4; **Fig. 2C**), consistent with the distance-based analysis. Instead, extra-TFIIIC sites were overrepresented among the chromatin states associated with repetitive elements including “Transcription elongation IV: low expression and repeats” (51 sites, 10.5%), “Repeats, intergenic, low expression introns” (85 sites, 17.5%), and “Repeat, RNA pseudogenes, H3K9me2” (192 sites, 39.5%) (permutation-based empirical P<0.001; **Fig. 2C**), consistent with CeRep3 repeat overrepresentation at extra-TFIIIC sites. Extra-TFIIIC sites were also overrepresented in “Enhancer II, intergenic” (40 sites, 8.2%; permutation-based empirical P<0.001; **Fig. 2C**) and “borders” between “active” and “regulated” chromatin domains known to harbor gene-distal transcription factor binding sites [4] (125 sites, 25%; permutation-based empirical P<0.001; **Fig. 2D**). However, extra-TFIIIC sites were devoid of histone modifications associated with active enhancers (H3K27ac), poised enhancers (H3K4me1), active promoters H3K4me3, or of Pol II enrichment (**Fig. 2E**). Pol III-bound TFIIIC sites were also not marked by H3K27ac, H3K4me1, or H3K4me3, but showed enrichment of Pol II, similar to previous observations [52]. Collectively, these results suggest that *C. elegans* extra-TFIIIC sites do not present features of active *cis-*regulatory elements for Pol II-dependent transcription.

### *C. elegans* extra-TFIIIC sites are densely clustered in the distal arms of autosomes

The lack of robust association with local regulatory features of Pol II and Pol III transcription led us to hypothesize that extra-TFIIIC sites were related to large-scale organization of chromosomes as in the case of yeasts [40,53]. To test this hypothesis, we examined the distribution of extra-TFIIIC sites in the genome. We observed that extra-TFIIIC sites were highly overrepresented in chromosome V (195 of the 504 sites, 39%; permutation-based empirical P<0.001), but strongly under-represented in the X chromosome (18 of the 504 sites, 4%; permutation-based empirical P<0.001; **Fig. 3A**). In contrast, Pol III-bound TFIIIC sites were highly over-represented in the X chromosome (202 of the 525 sites, 38%; permutation-based empirical P<0.001) consistent with tRNA gene overrepresentation in the X chromosome [23].

**FIGURE 3.**
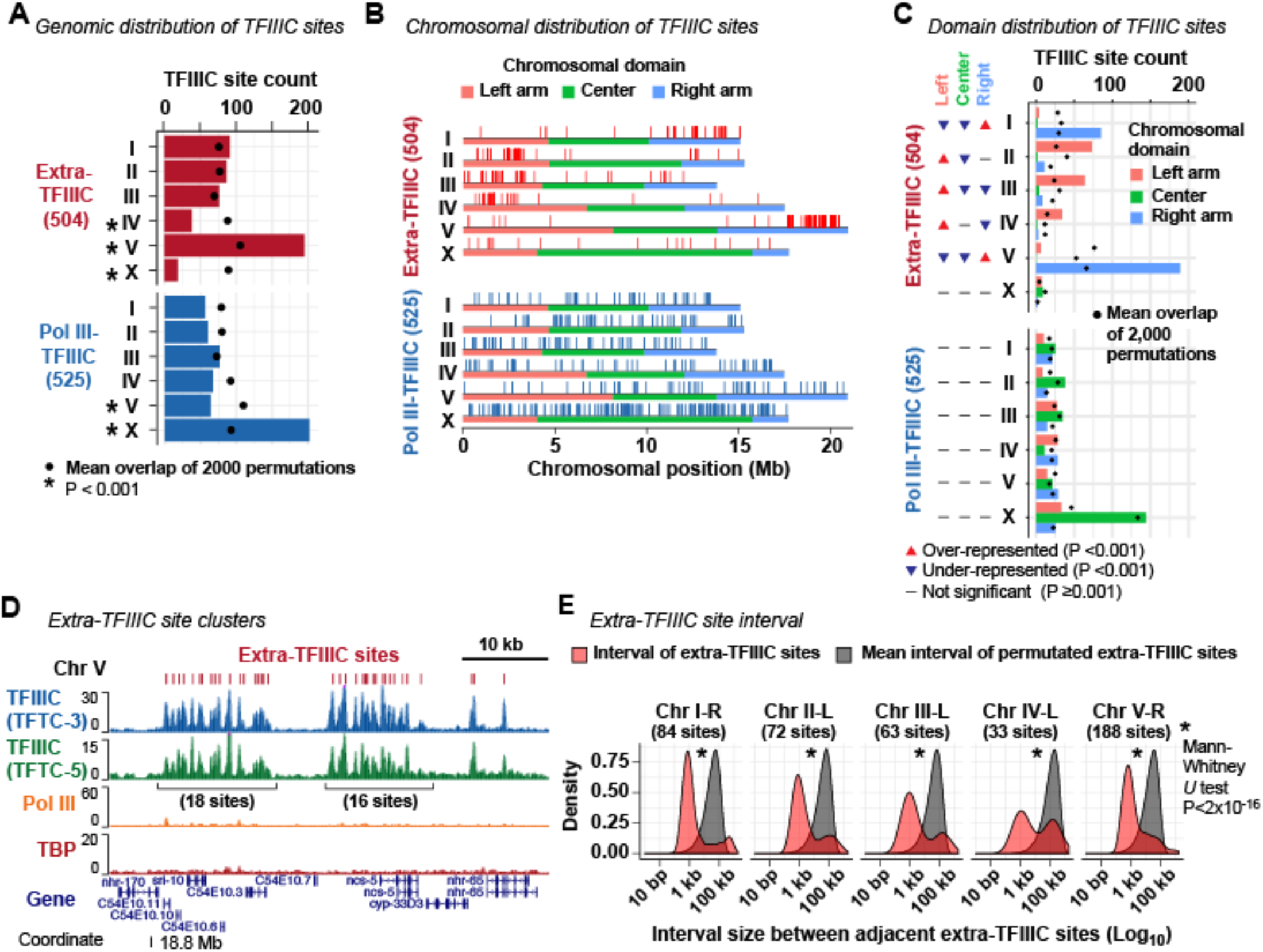
Extra-TFIIIC sites are clustered in autosomal arms. **(A)** Distribution of TFIIIC sites across chromosomes. Permutations of TFIIIC sites were performed across the genome. P, empirical P-value based on the 2000 permutations. **(B)** Distribution of TFIIIC sites within chromosomes. Horizontal color bars indicate chromosomal domains defined by Rockman and Kruglyak (2009). **(C)** Fraction of TFIIIC sites within chromosomal domains. Permutations of TFIIIC sites were performed within chromosomes. P, empirical P-value based on the 2000 permutations. **(D)** A representative genomic region harboring clusters of extra-TFIIIC sites. **(E)** Red, distribution of the interval size between two adjacent extra-TFIIIC sites. Gray, distribution of the mean of the interval sizes of permutated TFIIIC sites (2,000 means representing 2,000 permutations). Permutations were performed within chromosomal domains (i.e. center and left/right arms).

*C. elegans* autosomes can be subdivided into three domains of similar size (left arm, center, and right arm) based on repetitive element abundance, recombination rates, and chromatin organization [17,23,25]. We found that most extra-TFIIIC sites were located in autosome arms (486 of the 504 sites, 96%; **Fig. 3B**). In addition, extra-TFIIIC sites were overrepresented in only one of each autosome’s two arms that harbors the meiotic pairing centers [54] (overrepresented in the right arm of chromosome I; left arm of chromosome II; left arm of chromosome III; left arm of chromosome IV; right arm of chromosome V; permutation-based empirical P<0.001; **Fig. 3C**). Furthermore, within autosomal arms, extra-TFIIIC sites were locally densely clustered (**Fig. 3D**), with a median interval between neighboring extra-TFIIIC sites of 1207 bp (Mann Whitney *U*-test vs. within-arm permutations P=2×10^−16^; **Fig. 3E**). Among the autosome arms, the chromosome V right arm contained the largest number of extra-TFIIIC sites (188 sites) with extensive clusters (**Fig. 3B, D, E**). Thus, *C. elegans* extra-TFIIIC sites were fundamentally different from Pol III-bound TFIIIC sites in their genomic distribution and highly concentrated at specific locations within autosomal arm domains.

### *C. elegans* extra-TFIIIC sites intersperse H3K9me3-marked heterochromatin domains

The autosome arms in *C. elegans* exhibit high levels of H3K9me2 and H3K9me3, histone modifications associated with constitutive heterochromatin [16,17]. Furthermore, in each autosome, H3K9me2 and H3K9me3 signals are known to be stronger in one arm than the other [16,27]. We hypothesized that extra-TFIIIC sites were located near H3K9me2 or H3K9me3-marked regions because extra-TFIIIC sites have been implicated in heterochromatin insulation [55,56]. To test this hypothesis, we compared the locations of TFIIIC-bound sites with the locations of H3K9me2 and H3K9me3-enriched regions identified in early embryos [3]. Strikingly, the chromosome arms in which extra-TFIIIC sites were overrepresented coincided with the arms that exhibited strong H3K9me2 and H3K9me3 enrichment (**Fig. 4A, B**). However, at the local level, extra-TFIIIC sites did not reside in H3K9me3-enriched or H3K9me2-enriched regions (**Fig. 4C**), and were strongly underrepresented in H3K9me3-enriched regions (only 2% in H3K9me3-enriched regions; permutation-based P<0.001; **Fig. 4D**). Instead, extra-TFIIIC sites were located significantly closer to H3K9me2-enriched regions (median distance 3.1 kb) and H3K9me3-enriched regions (median distance 12.5 kb) compared with Pol III-bound TFIIIC sites (H3K9me2, median distance 34.8 kb, Mann Whitney *U*-test P<2×10^−16^; H3K9me3, median distance 50.1 kb, P<2×10^−16^) or extra-TFIIIC sites permutated within autosomal arms (H3K9me2, median distance 11.3 kb, Mann Whitney *U*-test P=1×10^−14^; H3K9me3, median distance 28.9 kb, P=7×10^−14^) (**Fig. 4E**). Our analysis thus revealed that *C. elegans* extra-TFIIIC sites were located close to, but not overlapped with, H3K9me2 and H3K9me3-enriched regions within autosomal arm domains.

**FIGURE 4.**
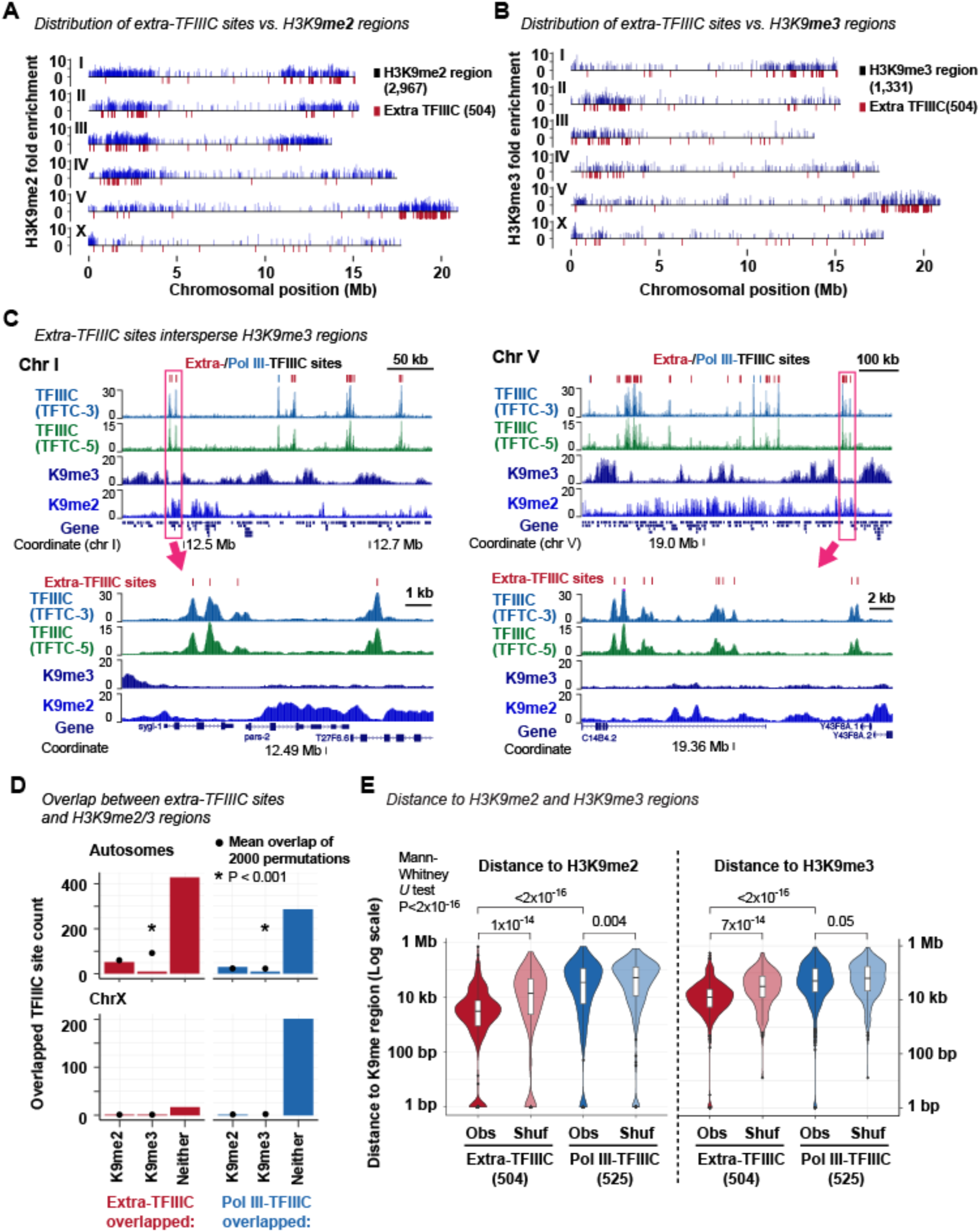
Extra-TFIIIC sites are located adjacent to H3K9me3-enriched regions. **(A)** Distribution of TFIIIC sites and H3K9me2-enriched regions. **(B)** Distribution of TFIIIC sites and H3K9me3-enriched regions. **(C)** (Top) Two representative genomic regions showing extra-TFIIIC sites interspersing H3K9me3-enriched regions. (Bottom) Genomic regions shown in rectangles in the top panels. **(D)** Fraction of TFIIIC sites overlapping H3K9me2/H3K9me3-enriched regions. Because the H3K9me2/H3K9me3-enriched regions are infrequent in the X chromosome, the X chromosome is plotted separately. Permutations of TFIIIC sites were performed within chromosomal domains (i.e. center and left/right arms). P, empirical P-value based on the 2000 permutations. **(E)** Distance between TFIIIC site center and H3K9me2- (left) or H3K9me3-enriched (left) regions. Obs, the observed distance. Shuf, distance between a permutated set of TFIIIC sites and H3K9me2-enriched regions. Permutation was performed within chromosomal domains.

### *C. elegans* extra-TFIIIC sites are located within nuclear membrane-associated domains

In *S. pombe* and *S. cerevisiae*, extra-TFIIIC sites are localized at the nuclear periphery and thought to regulate spatial organization of chromosomes [40,46]. In *C. elegans*, Pol III-transcribed genes including tRNA genes are associated with the nuclear pore component NPP-13 [49], similar to tRNA genes in *S. pombe* associated with the nuclear pores [57] (**Fig. 5A**). We hypothesized that extra-TFIIIC sites are associated with NPP-13, given the similarity of extra-TFIIIC sites to Pol III-transcribed genes. To test this hypothesis, we compared the locations of extra-TFIIIC sites with those of NPP-13-bound sites identified in mixed-stage embryos [49]. We observed that only 6 of the 504 extra-TFIIIC sites (1.2%) overlapped NPP-13-bound sites (**Fig. 5B**), in contrast to a large fraction of Pol III-bound TFIIIC sites (215 sites, 41%) that overlapped NPP-13-bound sites (permutation-based P<0.001; **Fig. 5B**). Thus, extra-TFIIIC sites were not likely to be associated with the nuclear pore.

**FIGURE 5.**
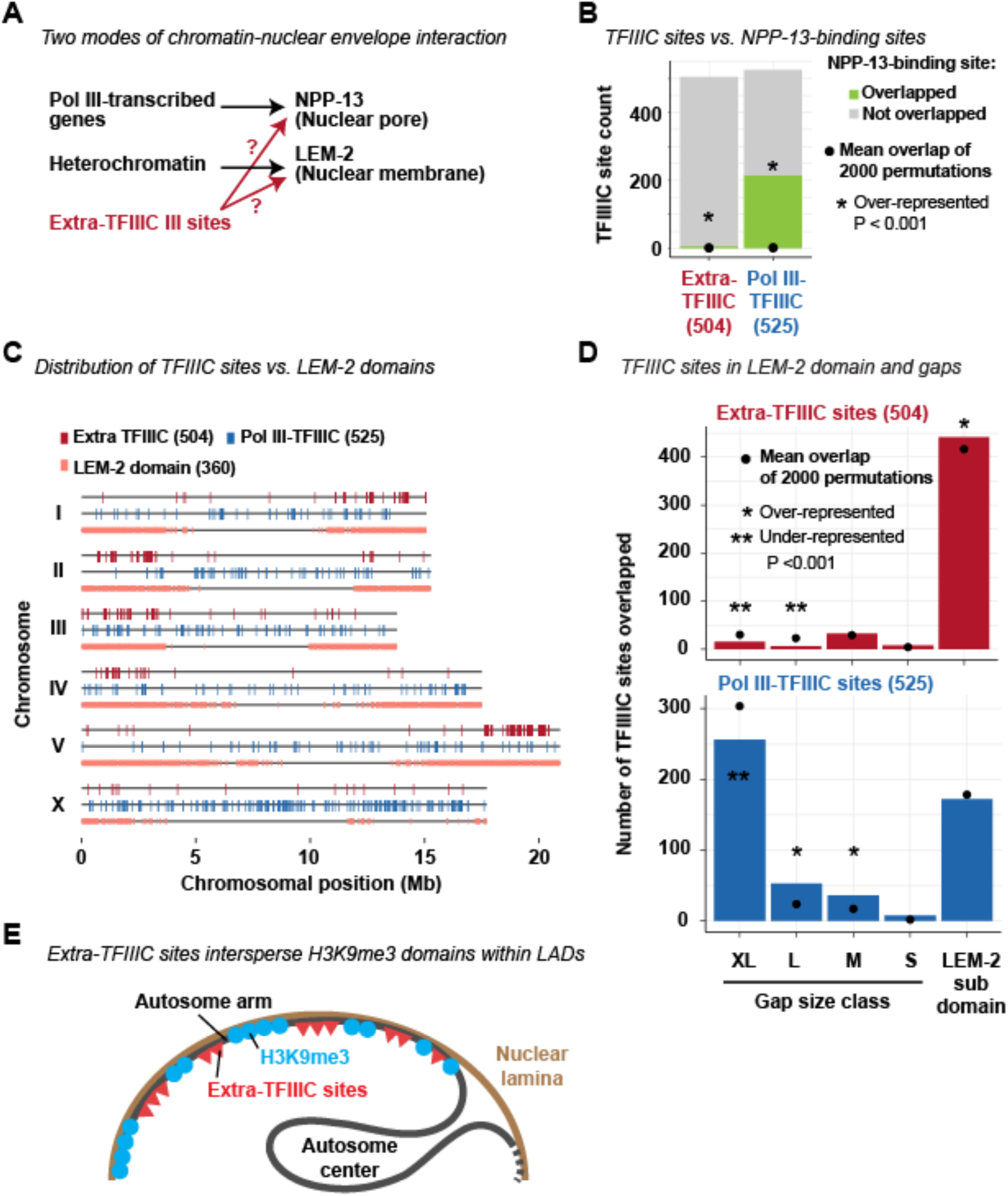
Extra-TFIIIC sites reside in LEM-2-associated domains. **(A)** Two hypothetical scenarios in which extra-TFIIIC sites associate with the nuclear periphery. **(B)** Fraction of TFIIIC sites overlapping 223 NPP-13-binding sites (±500 bp of NPP-13 site center) defined by Ikegami and Lieb (2013). Permutations of TFIIIC sites were performed within chromosomal domains (i.e. center and left/right arms). P, empirical P-value based on the 2000 permutations. **(C)** Distribution of TFIIIC sites and LEM-2 subdomains defined by Ikegami et al. (2010). **(D)** Fraction of TFIIIC sites overlapping LEM-2 subdomains and gaps of various sizes between LEM-2 subdomains. Permutations and P-value computation are performed as in **(B). (E)** Model for chromosomal organization where extra-TFIIIC sites are clustered in the autosome arms corresponding LEM-2 domains and intersperse H3K9me3-enriched regions.

Another mode of chromatin-nuclear envelope interactions in *C. elegans* is mediated by nuclear membrane-anchored, lamin-associated protein LEM-2 [28] (**Fig. 5A**). LEM-2 associates with the large genomic regions called “LEM-2 subdomains” that occupy 82% of the autosome arms [28]. Between LEM-2 subdomains are non-LEM-2-associated “gap” regions of various sizes [28]. We therefore investigated the locations of extra-TFIIIC sites with respect to those of LEM-2 subdomains and gaps (**Fig. 5C**). Strikingly, 441 of the 504 extra-TFIIIC sites (88%) were located within LEM-2 subdomains, demonstrating overrepresentation in a statistical test that accounted for the overrepresentation of extra-TFIIIC sites within autosome arms (permutation-based P<0.001, permutation performed within chromosomal center/arm domains) (**Fig. 5C, D**). In contrast, Pol III-bound TFIIIC sites were underrepresented within LEM-2 associated domains (permutation-based P<0.001, permutation performed within chromosomal domains). Together, our results suggest that *C. elegans* extra-TFIIIC sites are localized at the nuclear periphery and intersperse H3K9me3-marked heterochromatin regions (**Fig. 5E**).

## DISCUSSION

In this paper, we identified genomic sites bound by Transcription Factor IIIC (TFIIIC), the basal transcription factor for RNA Polymerase III (Pol III), that do not participate in Pol III-dependent transcription in *C. elegans*. Similar TFIIIC-bound sites devoid of Pol III-dependent transcription, termed extra-TFIIIC sites, have been reported in yeast [39,40], fly [41], mouse [42], and human [43]. Our data demonstrated that half of all TFIIIC-bound sites in *C. elegans* embryos lack Pol III binding, TBP binding, and nearby noncoding RNA genes, revealing pervasive extra-TFIIIC sites in the *C. elegans* genome.

Previous studies have suggested that extra-TFIIIC sites act as genomic insulators by blocking enhancer activity or heterochromatin spreading [40,56,58], or mediating three-dimensional genome interactions [41,58]. Some of the genomic and chromatin features of *C. elegans* extra-TFIIIC sites reported in this paper resemble characteristics of extra-TFIIIC sites participating in chromatin insulation. First, *C. elegans* extra-TFIIIC were densely clustered in *cis*, similar to clusters of TFIIIC-bound sites capable of insulating heterochromatin and enhancer activities in *S. pombe* and human cells [40,58]. Second, *C. elegans* extra-TFIIIC sites were located close to, but not within, H3K9me3-marked regions, similar to the observation that some extra-TFIIIC sites are located at the boundaries of heterochromatin [40,56,58]. Third, *C. elegans* extra-TFIIIC sites coincided with genomic regions known to be associated with nuclear membrane protein LEM-2 [28], similar to yeast extra-TFIIIC sites localized to the nuclear periphery [40,46] These observations raise the possibility that *C. elegans* extra-TFIIIC sites may act as a chromatin insulator at the nuclear periphery.

There are also differences between *C. elegans* extra-TFIIIC sites and those in other organisms. First, *C. elegans* extra-TFIIIC sites possess both the Box-A and Box-B motifs, unlike yeast and human extra-TFIIIC sites that only possess the Box-B motif [39,43]. Second, *C. elegans* extra-TFIIIC sites were neither located near gene promoters nor associated with chromatin features of Pol II-dependent regulatory regions, unlike human, fly, and mouse extra-TFIIIC sites that are located near regulatory elements for RNA Polymerase II (Pol II) transcription [41–43]. Third, *C. elegans* extra-TFIIIC sites are unrelated to CTCF binding, unlike human and mouse extra-TFIIIC sites that are located near CTCF binding sites [42,43], because the *C. elegans* genome does not encode CTCF [32]. Our study could thus offer an opportunity for a comparative analysis of extra-TFIIIC functions across eukaryotic species.

The molecular mechanisms underlying the chromatin organization of *C. elegans* autosomes remain poorly understood. Unlike the X chromosome organized into large self-interacting domains that have some features shared with topological associated domains (TADs) reported in other organisms, the autosomes do not present strong and pervasive self-interacting domains [22]. The condensin binding sites that could create boundaries between self-interacting domains in the X chromosome did not do so in the autosomes [14]. Several mechanisms for autosome chromatin organization have been proposed. These mechanisms include the antagonism between H3K36 methyltransferase MES-4 and H3K27 methyltransferase RPC2 that defines active versus repressed chromatin boundaries [4,34]; small chromatin loops emanating from the nuclear periphery that allow active transcription within heterochromatin domains [28]; and active retention of histone acetylase to euchromatin that prevents heterochromatin relocalization [33]. Our data that extra-TFIIIC sites are highly overrepresented in autosome arms and cluster in *cis* near the boundaries of H3K9me3-marked regions warrant future investigation of whether TFIIIC proteins participate in chromatin organization in *C. elegans* autosomes.

How strategies to demarcate chromatin domains have evolved in eukaryotes remain unclear. In vertebrates, CTCF has a central role in defining TAD boundaries and is essential for development [11,59,60]. In *D. melanogaster*, CTCF is essential but does not appear to define TAD boundaries, and instead acts as a barrier insulator [61–63]. In the non-bilaterian metazoans, some bilaterian animals (such as *C. elegans*), plants, and fungi, CTCF orthologs are absent [32,64]. In contrast to CTCF, the TFIIIC proteins are conserved across eukaryotes [65] and extra-TFIIIC sites have been reported in human, [43], mouse [42], fly [41], *C elegans* (this study), and yeast [39,40]. Whether extra-TFIIIC is an evolutionary conserved mechanism for demarcating chromatin domains in eukaryotes, including those lacking a CTCF ortholog, will be an interesting subject of future studies.

## CONCLUSIONS

We identified TFIIIC-bound sites that do not participate in RNA Polymerase III-dependent transcription in the *C. elegans* genome. These “extra-TFIIIC” sites were highly over-represented in the arm domains of the autosomes interacting with the nuclear lamina. Extra-TFIIIC sites formed dense clusters in *cis* near the outer boundaries of individual H3K9me3-marked heterochromatin regions. These genomic features of *C. elegans* extra-TFIIIC sites resemble extra-TFIIIC sites reported in other organisms that have activities in insulating heterochromatin. Our study warrants future investigation of whether TFIIIC proteins participate in heterochromatin insulation in *C. elegans*.

## METHODS

### ChIP-seq dataset

ChIP-seq of TFTC-3, TFTC-5, RPC-1, and TBP-1 was performed in chromatin extracts of the mixed-stage N2-strain embryos in duplicates and have been reported in our previous publication [49]. These data sets are available at Gene Expression Omnibus (GEO; http://www.ncbi.nlm.nih.gov/geo/) with accession numbers GSE28772 (TFTC-3 ChIP and input), GSE28773 (TFTC-5 ChIP and input), GSE28774 (RPC-1 ChIP and input), and GSE42714 (TBP-1 ChIP and input). ChIP-seq datasets for H3K4me3, H3K4me1, H3K27ac, H3K9me2, and H3K9me3, performed in chromatin extracts of early-stage N2-strain embryos [3], were downloaded from ENCODE website (https://www.encodeproject.org/comparative/chromatin/).

### Reference genome

The ce10 reference sequence was used throughout. The chromosomal domains (left arm, center, and the right arm) defined by recombination rates [24] was used.

### Gene annotation

The genomic coordinates and the types of *C. elegans* transcripts were downloaded from the WS264 annotation of WormMine. The WS264 genomic coordinates were transformed to the ce10 genomic coordinates using the liftOver function (version 343) with the default mapping parameter using the *ce11ToCe10*.*over*.*chain* chain file downloaded from the UCSC genome browser.

### TFIIIC site definition

MACS2 identified 1,658 TFTC-3-enriched sites. Of those, sites that had the TFTC-3 fold-enrichment (FE) score greater than 5, harbored TFTC-5-binding sites within 100 bp, and were located in the nuclear chromosomes were considered “high-confidence” TFIIIC-bound sites (1,029 TFIIIC sites). The “center” of each TFIIIC site was defined by the position of the base with the largest TFTC-3 FE score. Of the 1,029 TFIIIC sites, those with the maximum Pol III (RPC-1) FE greater than 20 within ±250 bp of the site center were defined as “Pol III-bound TFIIIC” sites (525) and the remaining sites were defined as “extra-TFIIIC” sites (504). The genomic coordinates for Pol III-bound TFIIIC sites and extra-TFIIIC sites are listed in **Additional file 1**.

### Heatmap

For the heatmaps around TFIIIC sites, a set of 20-bp windows with a 10-bp offset that covered a 2-kb region centered around the center of TFIIIC sites was generated for each site. For each window, the mean of fold-enrichment score was computed from replicate-combined input-normalized fold enrichment bedgraph files. The signals were visualized using the ggplot2’s *geom_raster* function (version 2.2.1) in R.

### Non-coding RNA genes

The genomic location and classification of non-coding RNA genes was described in the *Gene Annotation* section. For each TFIIIC site extended +/–100 bp from the site center, whether the region contained the transcription start site (TSS) of non-coding RNA genes was assessed using the Bedtools *intersect* function [66] (version 2.26.0).

### DNA motif analysis

To find DNA motifs *de novo*, 150-bp sequences centered around the center of the TFIIIC-bound sites were analyzed by MEME (v4.11.3) [67] with the following parameters: minimum motif size, 6 bp; maximum motif size, 12 bp; and the expected motif occurrence of zero or one per sequence (-mod zoops) and with the 1st-order Markov model (i.e. the dinucleotide frequency) derived from the ce10 genome sequence as the background.

### Genomic intersection and permutation

Unless otherwise noted, the overlap between the 1-bp center of each of the TFIIIC sites and genomic features of interest (with size ≥ 1 bp) was assessed using the Bedtools *intersect* function [66] (version 2.26.0). To estimate the probability of observing the overlap frequency by chance given the frequency, location, and size of the features of interest and TFIIIC sites, the TFIIIC sites were permutated using the Bedtools *shuffle* function (version 2.26.0). The TFIIIC sites were shuffled across the genome, or within the chromosomes, or within chromosomal domains in which they reside, as described in each analysis section. After each permutation, the permutated set of TFIIIC sites were assessed for the overlap with the features of interest. This permutation was repeated 2,000 times to assess the frequency at which the number of intersections for the permutated set of the TFIIIC sites was greater or less than the number of intersections for the observed TFIIIC sites. If none of the 2,000 permutations resulted in the number of overlaps greater or less than the observed number of overlaps, the observed degree of overlaps was considered overrepresented or underrepresented, respectively, with the empirical P-value cutoff of 0.001. The mean number of overlaps after 2,000 permutations was computed for visualization.

### Repetitive element analysis

The ce10 genomic coordinate and classification of repetitive elements, compiled as the “RepeatMasker” feature, were downloaded from the UCSC genome browser. The intersection between repetitive elements and TFIIIC sites was assessed as described in the *Genomic intersection and permutation* section. The permutation of TFIIIC sites was performed within the chromosomes in which they resided.

### Protein-coding gene distance

The genomic location of protein-coding genes was described in the *Gene Annotation* section. For each TFIIIC site, the absolute distance between the center of the TFIIIC site and the closest TSS of a protein-coding gene was obtained using the Bedtools *closest* function [66] (version 2.26.0). To assess the probability of observing such distance distribution by chance given the frequency and location of the TSSs and TFIIIC sites, the TFIIIC sites were permutated once using the Bedtools *shuffle* function (version 2.26.0) such that the TFIIIC sites were shuffled within the chromosomes, and the distance between the permutated TFIIIC sites and closest protein-coding gene TSS was obtained. Mann-Whitney U test, provided by the *wilcox*.*test* function in R, was used to assess the difference of the distribution of the TFIIIC-TSS distances between groups.

### TFIIIC sites in exons, introns, and 3′-UTRs

The genomic coordinates of exons, introns, and 3′-UTRs were extracted from WS264 gff3 file downloaded from wormbase. Exons, introns, and 3′-UTRs that belong to protein-coding genes (i.e. transcript type is “coding_transcript”) were processed. The genomic coordinates were transformed to the ce10 genomic coordinates as described above. The intersection between these features and TFIIIC sites was assessed as described in the *Genomic intersection and permutation* section. The permutation of TFIIIC sites was performed within the chromosomal domains (see *Reference genome)*.

### Chromatin state analysis

The chromatin state annotations and annotations of “active”, “regulated”, and “border” domains are reported previously [4]. The intersection between chromatin state annotations and TFIIIC sites was assessed as described in the *Genomic intersection and permutation* section. The permutation of TFIIIC sites was performed within the chromosomal domains (see *Reference genome)* to account for the difference of the chromatin state representation among different chromosomal domains.

### Chromosomal distribution of TFIIIC sites

The number of TFIIIC sites in each chromosome was assessed as described in the *Genomic intersection and permutation* section. The permutation of TFIIIC sites was performed across the genome. The number of TFIIIC sites in each chromosomal domain (see *Reference genome)* was assessed as described in the *Genomic intersection and permutation* section. The permutation of TFIIIC sites was performed within chromosomes.

### Extra-TFIIIC site interval

For each chromosomal domain, the genomic distance between every pair of two neighboring extra-TFIIIC sites (center-to-center distance) was computed in R. To estimate the degree of closeness between extra-TFIIIC sites only explained by the frequency of extra-TFIIIC sites within chromosomal domains, the extra-TFIIIC sites were permutated 2,000 times within chromosomal domains as described in the *Genomic intersection and permutation* section. In each of the 2,000 permutations, the genomic distance between two neighboring permutated extra-TFIIIC sites was computed, and the mean of the distances was computed. The distribution of the 2,000 means (by 2,000 permutations) was compared with the distribution of observed distribution of TFIIIC interval sizes by Mann-Whitney *U* test.

### Analysis of H3K9me2 and H3K9me3 regions

To define H3K9me2-enriched and H3K9me3-enriched regions, the genome was segmented into 1-kb windows, and the mean fold-enrichment score of H3K9me2 and H3K9me3 (see *ChIP-seq dataset*) was computed for each window using the Bedtools *map* function (version 2.26.0). Windows with the mean fold-enrichment score greater than 2.5 (1.5x standard deviation above mean for H3K9me2; and 1.3x standard deviation above mean for H3K9me3) were considered enriched for H3K9me2 or H3K9me3 and merged if located without a gap. This yielded 2,967 H3K9me2-enriched regions (mean size, 2.1 kb) and 1,331 H3K9me3-enriched regions (mean size, 4.9 kb).

The intersection between H3K9me2-enriched or H3K9me3-enriched regions and TFIIIC sites was assessed as described in the *Genomic intersection and permutation* section. The permutation of TFIIIC sites was performed within the chromosomal domains.

For each TFIIIC site, the absolute distance between the center of the TFIIIC site and the closest H3K9me2-enriched and H3K9me3-enriched regions was obtained using the Bedtools *closest* function [66] (version 2.26.0). To assess the probability of observing such distance distribution by chance given the frequency, location, and size of H3K9me2-enriched and H3K9me3-enriched regions and TFIIIC sites, the TFIIIC sites were permutated once using the Bedtools *shuffle* function (version 2.26.0). For the analysis of the distance to H3K9me2-enriched regions, this permutation was performed within the chromosomal domains. For the analysis of the distance to H3K9me3-enriched regions, permutation was performed with the chromosomal domains but excluding the H3K9me3-enriched regions themselves because the TFIIIC sites were strongly underrepresented in the H3K9me3-enriched regions.

### TFIIIC sites in NPP-13-binding sites

The 223 NPP-13-binding sites identified in mixed-stage N2-stage embryos are previously reported [49]. The genomic coordinates were converted to the ce10 genomic coordinates using the UCSCtools *liftOver* function (version 343). The intersection between NPP-13-binding sites (±500 bp of NPP-13 binding site center) and TFIIIC sites was assessed as described in the *Genomic intersection and permutation* section. The permutation of TFIIIC sites was performed within the chromosomal domains.

### TFIIIC sites in LEM-2 subdomain and gaps

The LEM-2 subdomains and gaps between LEM-2 subdomains identified in mixed-stage N2-stage embryos are previously reported [28]. The genomic coordinates were converted to the ce10 genomic coordinates using the UCSCtools *liftOver* function (version 343). The intersection between LEM-2 subdomains or gaps of variable size classes and TFIIIC sites was assessed as described in the *Genomic intersection and permutation* section. The permutation of TFIIIC sites was performed within the chromosomal domains.

## LIST OF ABBREVIATIONS

*C. elegans*: *Caenorhabditis elegans*
*D. melanogaster*: *Drosophila melanogaster*
*S. cerevisiae*: *Saccharomyces cerevisiae*
*S. pombe*: *Schizosaccharomyces pombe*
ChIP-seq: chromatin immunoprecipitation followed by sequencing
Pol II: RNA Polymerase II
Pol III: RNA Polymerase III
TBP: TATA-binding protein
TFIIIB: Transcription factor III B
TFIIIC: Transcription factor III C
TSS: Transcription start site(s)
H3K9me2: Histone H3 Lysine 9 dimethylation
H3K9me3: Histone H3 Lysine 9 trimethylation
H3K27ac: Histone H3 Lysine 27 acetylation
H3K4me1: Histone H3 Lysine 4 monomethylation
H3K4me3: Histone H3 Lysine 4 trimethylation
FE: Fold enrichment
TAD: Topologically Associated Domain

## DECLARATIONS

### ETHICS APPROVAL AND CONSENT TO PARTICIPATE

Not applicable.

### CONSENT FOR PUBLICATION

Not applicable.

### AVAILABILITY OF DATA AND MATERIALS

The datasets supporting the conclusions of this article are available in Gene Expression Omnibus with accession number GSE28772, GSE28773, GSE28774, and GSE42714 at https://www.ncbi.nlm.nih.gov/geo/. The datasets supporting the conclusions of this article are also included within the article (**Additional file 1**).

## AUTHOR CONTRIBUTIONS

K.I. conceived the study. K.I., A.S., A.L., and V.B. analyzed the data. K.I., A.S., A.L., and V.B. wrote the manuscript.

## ACKNOWLEDGEMENTS

We thank Jason D. Lieb, Sevinc Ercan, and Sebastian Pott for discussion.

